# Akt/PKB enhances non-canonical Wnt signals by compartmentalizing β-Catenin

**DOI:** 10.1101/149351

**Authors:** Nicolas Aznar, Nina Sun, Ying Dunkel, Jason Ear, Matthew D. Buschman, Pradipta Ghosh

**Author notes:** **Current address**: Department of Cancer Cell Plasticity, Centre de Recherche en Cancérologie de Lyon (CCRL), France. To whom correspondence should be addressed: N.A and P.G, **Nicolas Aznar, Ph.D**; Department of Medicine, University of California, San Diego School of Medicine, La Jolla, California 92093-0651., **Pradipta Ghosh, M.D**; Professor, Departments of Medicine and Cellular and Molecular Medicine, University of California, San Diego School of Medicine; George E. Palade Laboratories for Cellular and Molecular Medicine, 9500 Gilman Drive, Room 333; La Jolla, California 92093-0651.

## Abstract

Cellular proliferation is antagonistically regulated by canonical and non-canonical Wnt signals; their dysbalance triggers cancers. It is widely believed that the PI3-K→ Akt pathway enhances canonical Wnt signals by affecting transcriptional activity and stability of β-catenin. Here we demonstrate that the PI3-K→Akt pathway also enhances non-canonical Wnt signals by compartmentalizing β-catenin. By phosphorylating the phosphoinositide(PI)-binding domain of a multimodular signal transducer, Daple, Akt abolishes Daple’s ability to bind PI3-P-enriched endosomes that engage dynein motor complex for long-distance trafficking of β-catenin/E-cadherin complexes to pericentriolar recycling endosomes (PCREs). Phosphorylation compartmentalizes Daple/β-catenin/E-cadherin complexes to cell-cell contact sites, enhances non-canonical Wnt signals, and thereby, suppresses colony growth. Dephosphorylation compartmentalizes β-catenin on PCREs, a specialized compartment for prolonged unopposed canonical Wnt signaling, and enhances colony growth. Cancer-associated Daple mutants that are insensitive to Akt mimic a constitutively dephosphorylated state. This work not only identifies Daple as a platform for crosstalk between Akt and the non-canonical Wnt pathway, but also reveals the impact of such crosstalk during cancer initiation and progression.

## Introduction

The Wnt signaling pathway plays a crucial role in embryonic development, in tissue regeneration and in many other cellular processes including cell fate, adhesion, polarity, migration, and proliferation. Dysregulated expression of components within the Wnt pathway triggers many diseases, and most importantly, heralds cancer (Klaus & Birchmeier, 2008). The well-characterized canonical Wnt signaling pathway enhances the stability, nuclear localization and activity of β-catenin, and the downstream activation of genes targeted by the TCF/LEF transcription machinery. This canonical Wnt pathway is antagonized by a non-canonical Wnt signaling paradigm (Ishitani et al, 2003; Olson & Gibo, 1998; Torres et al, 1996), although it is unclear how this occurs. The non-canonical Wnt pathway induces the elevation of intracellular Ca^2+^ and activation of the small G proteins RhoA and Rac1 (Kuhl et al, 2000; Niehrs, 2001; Winklbauer et al, 2001), which regulate polarized cell movements and the planar polarity of epithelial cells (Kuhl et al, 2000; Mayor & Theveneau, 2014; Sheldahl et al, 1999). Dysregulation of the non-canonical Wnt pathway is widely believed to drive cancer via a two-faceted mechanism (McDonald & Silver, 2009)--**1)** In the normal epithelium, non-canonical Wnt signaling is protective; it suppresses tumorigenesis by antagonizing the canonical β-catenin/TCF/LEF pathway due in part by compartmentalizing β-catenin at the cell membrane to prevent its transcriptional activity (Bernard et al, 2008); how such compartmentalization is accomplished remains unknown. What is well-accepted though, is that inhibition of non-canonical Wnt signaling fuels neoplastic transformation (Grumolato et al, 2010; Ishitani et al, 2003; Medrek et al, 2009); **2)** In transformed cells, however, non-canonical Wnt signaling enhances cancer invasion/metastasis by activation of Rac1 and remodeling of the actin cytoskeleton (Yamamoto et al, 2009) and by upregulating CamKII and PKC (Dissanayake et al, 2007; Weeraratna et al, 2002). Little is known as to how non-canonical Wnt signaling switches from being a protective pathway to one that enhances cancer progression.

We recently demonstrated that Daple, a multi-modular signal transducer is an enhancer of non-canonical Wnt signaling downstream of Frizzled receptors (FZDRs) (Aznar et al, 2015). Mechanistically, Daple serves as a novel guanine-nucleotide exchange factor (GEF) for heterotrimeric (henceforth trimeric) G protein, Gαi, and directly binds FZD receptors. By binding both FZD and G protein, Daple and enables ligand-activated receptors to recruit and activate Gi, and trigger non-canonical Wnt signaling (Aznar et al, 2015). Activation of Gαi dictates several closely intertwined spatial and temporal aspects of post-receptor signaling events, including, reduction of cellular levels of cAMP and β-catenin, and release of ‘free’ Gβγ from heterotrimers, which activates non-canonical Wnt signaling pathways involved in cell motility (e.g., PI3K/Akt and Rac1). Consequently, Daple suppresses β-catenin/TCF/LEF transcription program, oncogenic transformation, anchorage independent growth and anchorage-dependent colony formation; these properties of Daple mirror the key anti-growth and anti-transformation phenotypes that define a tumor suppressor / anti-oncogene (Cooper, 2000). Daple also enhances all phenotypes that have been previously attributed to the prometastatic role of non-canonical Wnt signaling (Minami et al, 2010), e.g., induction of actin stress fibers, increase in 2D-cell migration after wounding, 3D-invasion through basement membrane proteins, and upregulation of genes that trigger EMT. Thus, the FZD-Daple-G protein signaling axis serves as a double-edged sword-- it functions as an tumor suppressor in normal epithelium, but acts as an oncogene in transformed cells to enhance tumor invasion and metastasis (Aznar et al, 2015).

Here we show that Daple is a downstream target of Akt kinase. Findings presented here not only reveal the cellular consequences of this phosphoevent, but also provide mechanistic insights into how aberrations, e.g., mutations, copy no. variations (CNVs), etc. in the tumor suppressor gene, Daple and the oncogene Akt, and their complex interplay may cooperatively affect β-catenin signaling during cancer initiation and progression.

## Results and Discussion

### Intracellular pool of Daple localizes predominantly to pericentriolar recycling endosomes (PCREs)

We previously described that Daple is a cytosolic protein that is recruited to the plasma membrane (PM) in cells responding to Wnt5A, where it co-localizes with FZD7 receptor (Aznar et al, 2015). Although localization to the PM is ligand-dependent and short-lived, we consistently observed the presence of an additional intracellular perinuclear pool of myc-tagged Daple by confocal immunofluorescence which appeared as clusters of vesicles (arrows; **Figure 1A**). A similar perinuclear pool was also seen when endogenous Daple was detected with a previously validated antibody (Aznar et al, 2015) in HeLa (**Figure 1B**), HEK and DLD1 (**Figure S1A**) cells; in some cell lines the cluster of vesicles was more compact than others. This perinuclear localization was preserved when we expressed mutant Daple constructs that are specifically defective in binding/activating Gαi proteins [F1675A, lacks a functional G protein binding and activating motif; (Aznar et al, 2015)] or binding Dvl [ΔPBM; lacks the C-terminally located PDZ-binding motif; (Oshita et al, 2003)] (**Figure S1B**), indicating that these functional interactions are not essential for Daple’s localization. Because perinuclear localization of Daple did not change during Wnt5A stimulation (**Figure 1A**), and was independent of Daple’s ability to bind or activate G proteins or bind Dvl (**Figure S1B**), this localization was deemed constitutive, and hence, all subsequent studies were carried out at steady-state in the presence of 10% serum.

**Figure 1:**
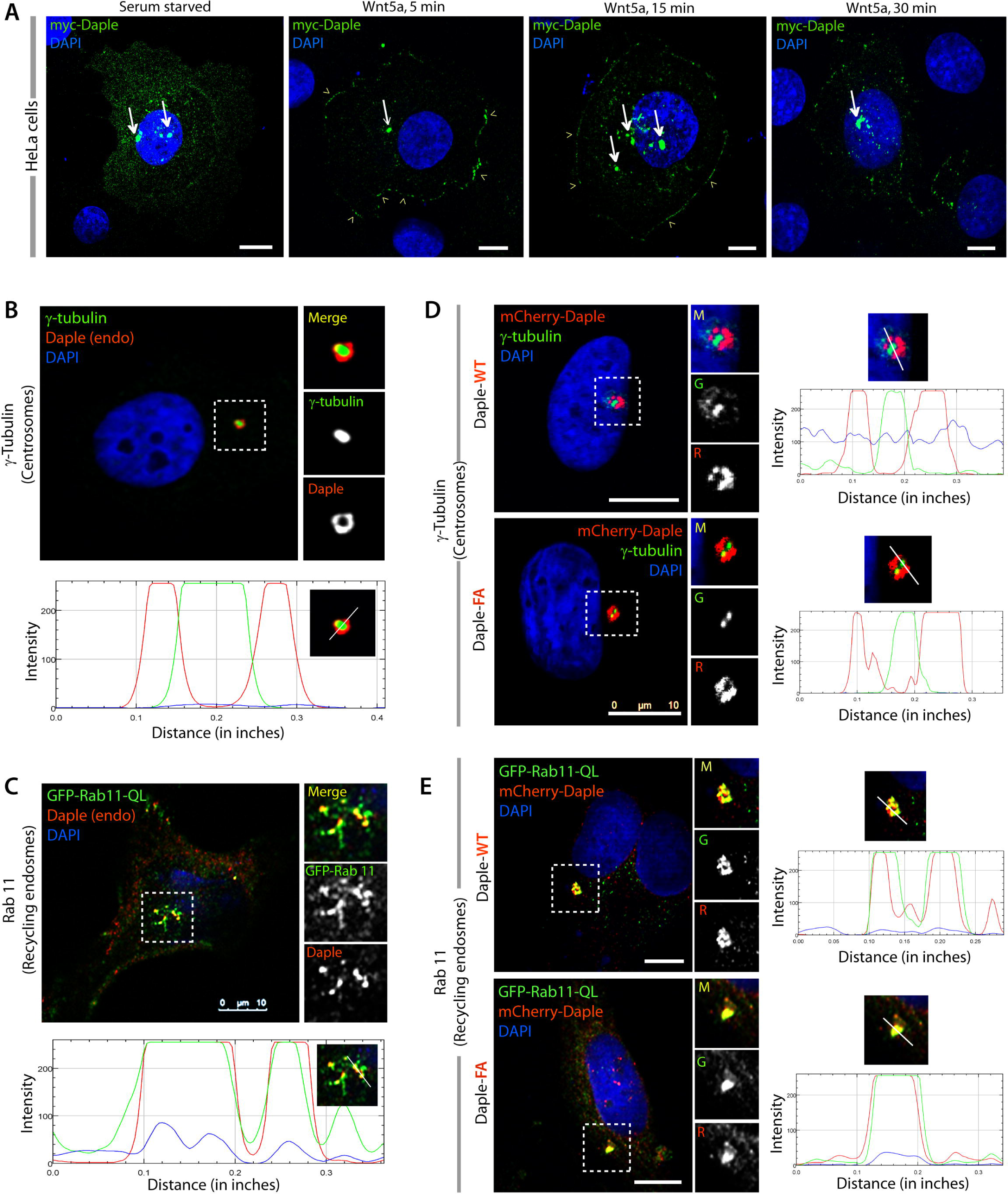
Daple localizes to peri-centriolar recycling endosomes (PCRE). (**A**) Serum-starved HeLa cells expressing myc-Daple WT were stimulated with Wnt5a for various time points, fixed, stained and analyzed for localization of myc-Daple (Green) by confocal microscopy. Myc-Daple localizes predominantly within an intracellular perinuclear pool (arrows) at steady-state. Upon Wnt5a stimulation, Daple transiently localizes to the plasma membrane (PM) (arrowheads; 15 min). (**B**) HeLa cells were fixed and stained for endogenous Daple (red), γ-tubulin (green) and DAPI (blue; nuclei) and analyzed by confocal microscopy. RGB plot, generated using ImageJ at the bottom shows how red pixels (Daple) flank green pixels (γ-tubulin; a centrosomal marker). (**C**) HeLa cells expressing GFP-Rab11-QL (a marker of recycling endosomes) were fixed and stained for endogenous Daple (red) and DAPI (blue; nuclei) and analyzed by confocal microscopy. RGB plot shows the degree of colocalization of red and green pixels. (**D, E**) HeLa cells transfected with either wild-type (WT) or GEF-deficient (FA) mCherry-Daple constructs were analyzed by confocal microscopy for their co-localization with either γ-tubulin (**D**) or GFP-Rab11-QL (**E**). RGB plots show that mCherry-Daple colocalized with Rab11 and around the centrosome regardless of a functional GEF motif.

To identify the nature of this perinuclear compartment, we performed immunofluorescence studies in Hela cells to look for colocalization between Daple and a variety of organelle markers (**Figure 1B-C; S1C-E**). Both endogenous and exogenously-expressed Daple localized around the centrosome (marked by γ-tubulin; **Figure 1B**), at the point of convergence of the stabilized acetylated microtubules (**Figure S1C**), and colocalized with the Rab GTPase, Rab11; the latter is a known coordinator of trafficking out of the pericentriolar recycling compartment to the PM (Grant & Donaldson, 2009; Ullrich et al, 1996) (**Figure 1C**). No colocalization was noted with early or late endosome markers, EEA1 and CD63, respectively (**Figure S1D**), and only partial colocalization was noted with autophagosome marker, LC3 (**Figure S1E**); the latter is consistent with the fact that autophagosome membranes coalesce with Rab11-containing recycling endosomes (Longatti & Tooze, 2012; Puri et al, 2013; Szatmari et al, 2014). Taken together, these findings pinpoint Daple’s localization to Rab11-positive pericentriolar recycling endosomes (PCREs). Furthermore, we confirmed that Daple’s ability to localize to PCREs or the appearance of the PCRE compartment was not impacted by Daple’s G protein modulatory functions because both WT and a GEF-deficient Daple-FA showed similar distributions around the centrosome (**Figure 1D**) and colocalized with Rab11-positive vesicles (**Figure 1E**). We conclude that a significant pool of Daple localizes to the PCREs, and that such localization is independent of ligand stimulation or its G protein modulatory function.

### Identification of putative phosphoinositide-binding domain and Akt phosphorylation site in Daple that is mutated in cancers

To determine which domain is important for the localization of Daple to PCREs, we generated two truncated myc-Daple constructs (**Figure 2A**) and analyzed their localization in HeLa cells by confocal immunofluorescence. Full length Daple WT or a short Daple construct corresponding to only a region of the coiled coil domain (aa 1194-1491) localized to the perinuclear compartment, whereas expression of the C-terminus alone (aa 1476-2028) localized predominantly at the PM and homogenously in the cytoplasm; no discernible enrichment was noted in any perinuclear compartment (**Figure 2A**). These findings suggest that a region within aa 1194-1491 may be sufficient and required for targeting Daple to PCREs.

**Figure 2:**
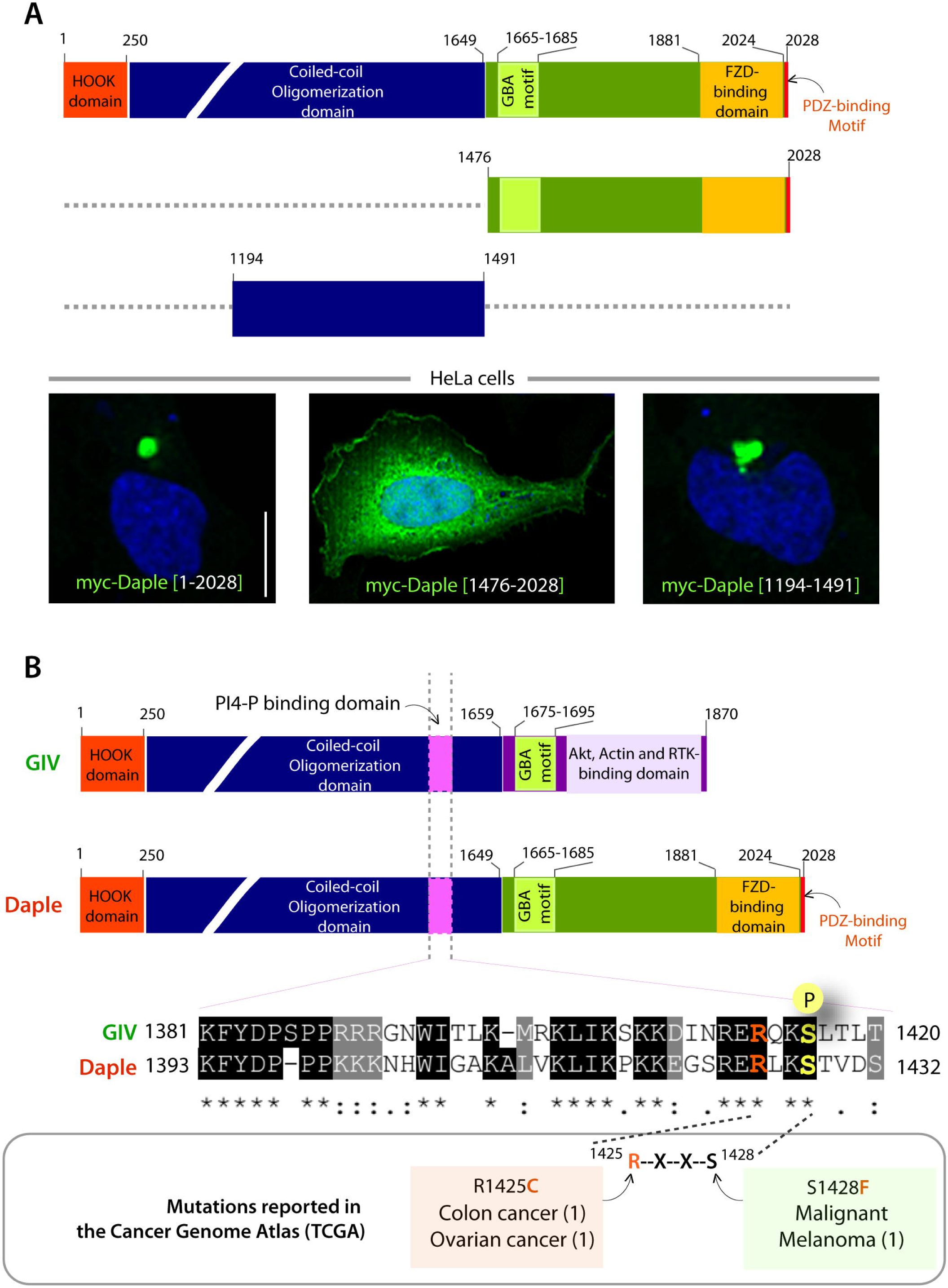
Identification of a putative lipid binding domain within Daple. (**A**) (Top) schematic showing the various functional modules of full length Daple and other truncated constructs used to identify domains regulating its subcellular localization. From the N- to the C-terminus the domains are– a predicted Hook-domain (red), which binds microtubules; a long coiled-coil domain (blue) assists in homo/oligomerization (Oshita et al, 2003); an evolutionarily conserved GEF motif (green) which binds and activates Gai (Aznar et al, 2015) and releases ‘free’ Gβγ. The C-terminal ~ 150 aa of Daple (orange) also has key domains that enable Daple to bind FZD7 Receptor (Aznar et al, 2015), and bind DVL (Oshita et al, 2003). (Bottom) HeLa cells transfected with either myc-Daple full length (aa 1-2028) or myc-Daple short constructs (aa 1476-2028 and aa 1194-1491) were analyzed for localization of myc-Daple (Green) and DAPI (blue; nuclei). (**B**) Sequence analysis revealed a motif in Daple similar to the PI4-P binding motif previously described in GIV (Enomoto et al, 2005). A previously described Akt phosphorylation site within GIV, S1416 is conserved on Daple (S1428) within a consensus sequence (RXXS) predicted to be phosphorylated by Akt. Schematic also lists mutations within or flanking the putative Akt phosphorylation site of Daple that have been reported in The Cancer Genome Atlas (TCGA) database. Numbers in parenthesis = total events.

A closer assessment of the sequence within the region between 1194-1491 revealed that Daple has a putative phosphoinositide(PI)-binding domain, much like its closely related paralogue, GIV/Girdin. Daple and GIV are members of the CCDC88 family; they share high degree of homology in their N-terminal Hook and coiled-coil domains, but are highly divergent in their C-terminal domains. Within the homologous N-terminal region, GIV has been shown to contain a PI-binding domain (aa 1381-1420), which specifically binds phosphoinisitol 4-phosphate at the PM and on Golgi membranes (PI4-P) (Enomoto et al, 2005). This GIV-PI4-P interaction is regulated by a single phosphoevent, i.e., phosphorylation of GIV at Ser(S)1416 by Akt dissociates the interaction (Enomoto et al, 2005; Ghosh et al, 2008). Alignment of Daple (aa 1194-1491) with the corresponding region on GIV revealed the presence of an evolutionarily conserved putative PI-binding domain containing a Akt consensus site (^1425^RXXS^1428^) (**Figure 2B; S2A**). Several high-throughput LCMS studies conducted in a variety of cell types using Akt phospho-substrate antibodies (Cell Signaling Inc.) had previously reported that S1428 within this consensus on Daple is phosphorylated in cells (**Figure S3**). Multiple kinase prediction programs (Scansite Motif Scan, MIT; NetPhos 2.0, Denmark; KinasePhos, Taiwan; Phospho-Motif Finder, HPRD) were also in agreement that Akt is predicted to phosphorylate Daple at Ser1428. These findings raised 3 possibilities: *1)* that like GIV, Daple may bind PI (lipid) via this putative PI-binding motif; *2)* Akt may phosphorylate the conserved S1428 within the PI-binding motif on Daple; and *3)* that this phosphoevent may regulate Daple’s ability to bind PI.

Furthermore, because Daple is a key regulator of tumor initiation and progression (Aznar et al, 2015), we looked for clues into how perturbation of Daple’s putative PI-binding domain affects cancer progression. A search of the comprehensive catalog of somatic mutations (COSMIC) database curated by genome sequencing in the Cancer Genome Atlas (TCGA) revealed that the sequence within the Akt-consensus site is recurrently mutated (**Figure 2B; S2B-C**). Of those, two mutations were of utmost interest to us : Ser(S)1428>Phe(F) [S1428F; which should render Daple non-phosphorylatable] and Arg(R)1425>Cys(C) [R1425C ; henceforth, RC], which is predicted to disrupt the Akt consensus site, and hence also predicted to be non-phosphorylatable. These revelations suggest that the newly identified putative PI-binding and Akt phosphorylation sites on Daple could be key regulators of the newly identified intracellular localization to PCREs, the previously defined Daple’s functions in cells and its role in tumor initiation and progression (Aznar et al, 2015).

### Phopshorylation of the phosphoinositide-binding domain by Akt regulates Daple’s ability to bind PI3-P

First, we wanted to confirm if Daple is indeed a substrate of Akt kinase. In vitro kinase assays confirmed that recombinant Akt kinase phosphorylated bacterially expressed His-tagged Daple-WT protein (aa 1194-1491), but not the non-phosphorylatable S1428A mutant [SA; in which Ser1428 is mutated to Ala], demonstrating that Akt can phosphorylate Daple in vitro at S1428. Disruption of the ^1425^RXXS^1428^ consensus in the tumor-associated His-Daple-RC mutant also abolished phosphorylation, demonstrating that the consensus sequence is essential for Akt-dependent phosphorylation *in vitro* (**Figure 3A**). These findings also held true in cells because when Daple was immunoprecipitated from cells with an anti-Akt phosphosubstrate, only Daple-WT, but not SA or RC mutants could be detected in the immunoprecipitates (**Figure 3B**). These findings demonstrate that- 1) Daple is phosphorylated at S1428; 2) the site is phosphorylated by Akt; and 3) that S1428 is the only site on Daple that is phosphorylated by Akt, much like the corresponding S1416 in GIV (Enomoto et al, 2005).

**Figure 3:**
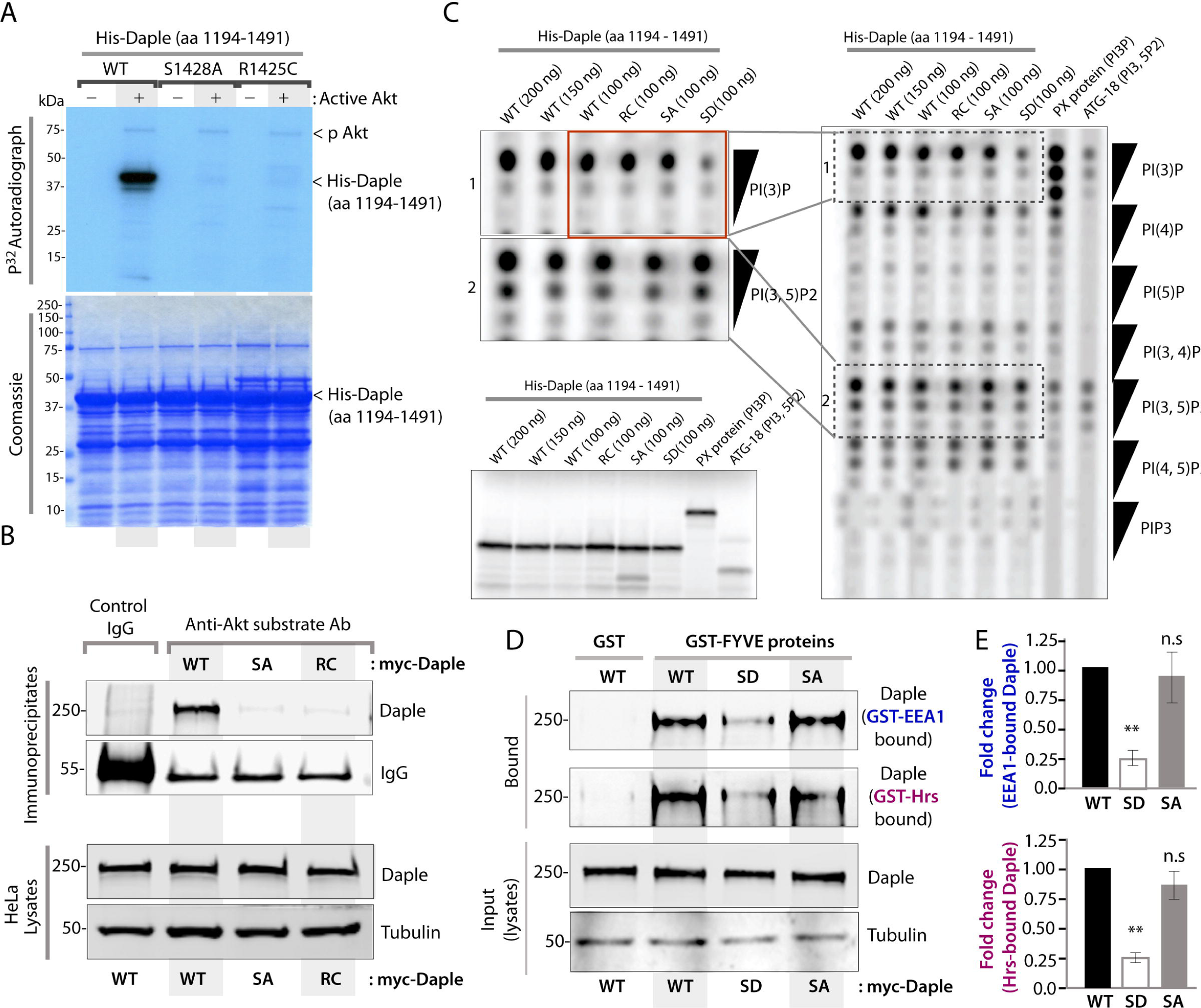
Daple binds PtdIns(3)P and PtdIns(3,5)P2; phosphorylation by Akt on Ser1428 regulates Daple’s ability to bind PI3-P. (**A**) *In vitro* kinase assays were carried out using recombinant Akt and bacterially expressed and purified His-Daple (aa 1194-1491) WT or the non-phosphorylatable (S1428A) or the cancer-associated (R1425C) proteins and γ-32P [ATP]. Phosphoproteins were analyzed by SDS-PAGE followed by autoradiography (top). Equal loading of substrate proteins was confirmed by staining the gel with Coomassie blue (bottom). (**B**) Daple is phosphorylated on Ser1428 by Akt in cells. Daple was immunoprecipitated from equal aliquots of lysates of HeLa cells expressing myc-Daple [WT, S1428A or R1425C mutants] using anti-Akt-substrate antibody or control IgG. Immune complexes (top) were analyzed by immunoblotting (IB) for Daple. (**C**) Daple binds PI3P and PI3,5-P2. The ability of His-Daple WT and either phosphomimic (SD) or non-phosphorylatable (SA and RC) His-Daple mutants to bind to a variety of phophoinositides was analysed using a protein–lipid-binding assay. His-PX and His-ATG-18 proteins, which are known to bind PtdIns(3)P and PtdIns(3,5)P2 respectively were used as a positive control. (Bottom left) Amount of in vitro-translated His-Daple constructs and controls used were analyzed by autoradiography. (**D, E**) Daple binds to PtdIns(3)P-enriched membranes. Crude membranes isolated from HeLa cells expressing Daple-WT, SD (phosphomimic) or SA (non-phosphorylatable) mutants were incubated with GST-FYVE domains of EEA1 or Hrs. Bound membranes were analysed for Daple by immunoblotting (**D**). Bar graphs (**E**) display the fold change in EEA1 (top) or Hrs (bottom)-bound Daple in **D**. Error bars representing mean ± S.D of three independent experiments.

Next we asked if Daple’s putative PI-binding motif is functional, i.e., capable of binding lipids, and if so, how this function may be impacted by the newly identified phosphoevent. To answer these questions, we generated an additional mutant, S1428> Asp(D), to mimic a constitutively phosphorylated state. Protein-lipid binding assays, as determined by lipid dot blots carried out using *in vitro* translated His-Daple protein revealed that Daple primarily binds to two types of lipids, PI3-P and PI3,5-P2 (**Figure 3C**); additional weaker interactions were seen also with PI4-P >> PI4,5P2, in decreasing order for affinity. No binding was seen for PIP3, PI3,4-P2, and PI5-P. Binding of Daple to PI3,5-P2 remained unchanged across WT and mutants. In the case of PI3-P, Daple-WT and the non-phosphorylatable SA and RC mutants bound equally, but binding was specifically reduced for the phosphomimicking Daple-SD mutant (**Figure 3C**). These findings indicated that Daple binds PI3-P and perhaps also PI3,5-P2 *in vitro*, but phosphorylation at S1428 selectively reduce the Daple-PI3-P interaction, without perturbing the Daple-PI3,5-P2 interaction.

To determine if these findings hold true in cells, we asked if Daple-WT and mutants associate with PI3-P enriched membranes isolated from cells using detergent-free homogenates of crude membrane fractions and previously validated PI3-P binding probes, GST-2xFYVE domains of EEA1 and Hrs (Gillooly et al, 2000). Daple-WT and SA were detected in these PI3-P enriched membranes, whereas binding was reduced in the case of Daple-SD (**Figure 3D-E**). These findings demonstrate that Daple binds PI3-P, and that binding is reduced when S1428 is phosphorylated. We conclude that phosphorylation of Daple by Akt within its lipid binding domain inhibits its affinity to PI3-P-enriched membranes. It is noteworthy that despite the homology, Daple’s PI-binding profile was strikingly different from that of GIV; the latter preferentially binds PI4-P (Enomoto et al, 2005), but not PI3-P. These differential preferences, however, are consistent with the differential patterns of their intracellular pools-- GIV primarily localizes to Golgi membranes (Le-Niculescu et al, 2005) that are enriched in PI4-P (Di Paolo & De Camilli, 2006), whereas Daple localizes to endosomes, which are enriched in PI3-P (Di Paolo & De Camilli, 2006; Haucke & Di Paolo, 2007). Although the determinants of such divergent PI-preferences of GIV and Daple is unclear, the parallelism between the two family members is striking; both GIV and Daple enhance Akt signaling (Anai et al, 2005; Aznar et al, 2015) and both are also substrates of Akt kinase [this work and (Enomoto et al, 2005)]; Akt phosphorylates both their PI-binding motifs, at one specific residue within the entire protein, and that single phosphoevent is sufficient to disrupt protein-lipid binding in both cases.

### Phosphoregulation of Daple’s PI-binding domain by Akt regulates the co-distribution of Daple and β-catenin at cell-cell junctions and PCREs

Next we asked what cargo proteins may be shuttled via the Daple-labeled PCREs. We previously showed that Daple is essential for the maintenance of low cytosolic levels of β-catenin; in cells without Daple, β-catenin is stabilized and levels of this protein rise (Aznar et al, 2015). Immunofluorescence studies in extensively validated Hela cell lines (Aznar et al, 2015) with (control; shLuc) of without (shDaple) Daple revealed that the distribution pattern of β-catenin in both cells is similar, i.e., at the PM and in clusters of perinuclear vesicles (**Figure 4A; left**). The perinuclear vesicular cluster, however, was more diffuse and the intensity of signal was increased both at the PM and in the perinuclear compartment in Daple-depleted cells, consistent with the previously reported ~3-fold elevated levels of β-catenin in these cells (Aznar et al, 2015). Because β-catenin/E-cadherin macrocomplexes are known to be shuttled to and from the PM via Rab11 positive PCREs (Horgan et al, 2010; Kam & Quaranta, 2009; Lock & Stow, 2005), we hypothesized that Daple may colocalize with these macrocomplexes and impact their trafficking between the PM and the PCREs.

**Figure 4:**
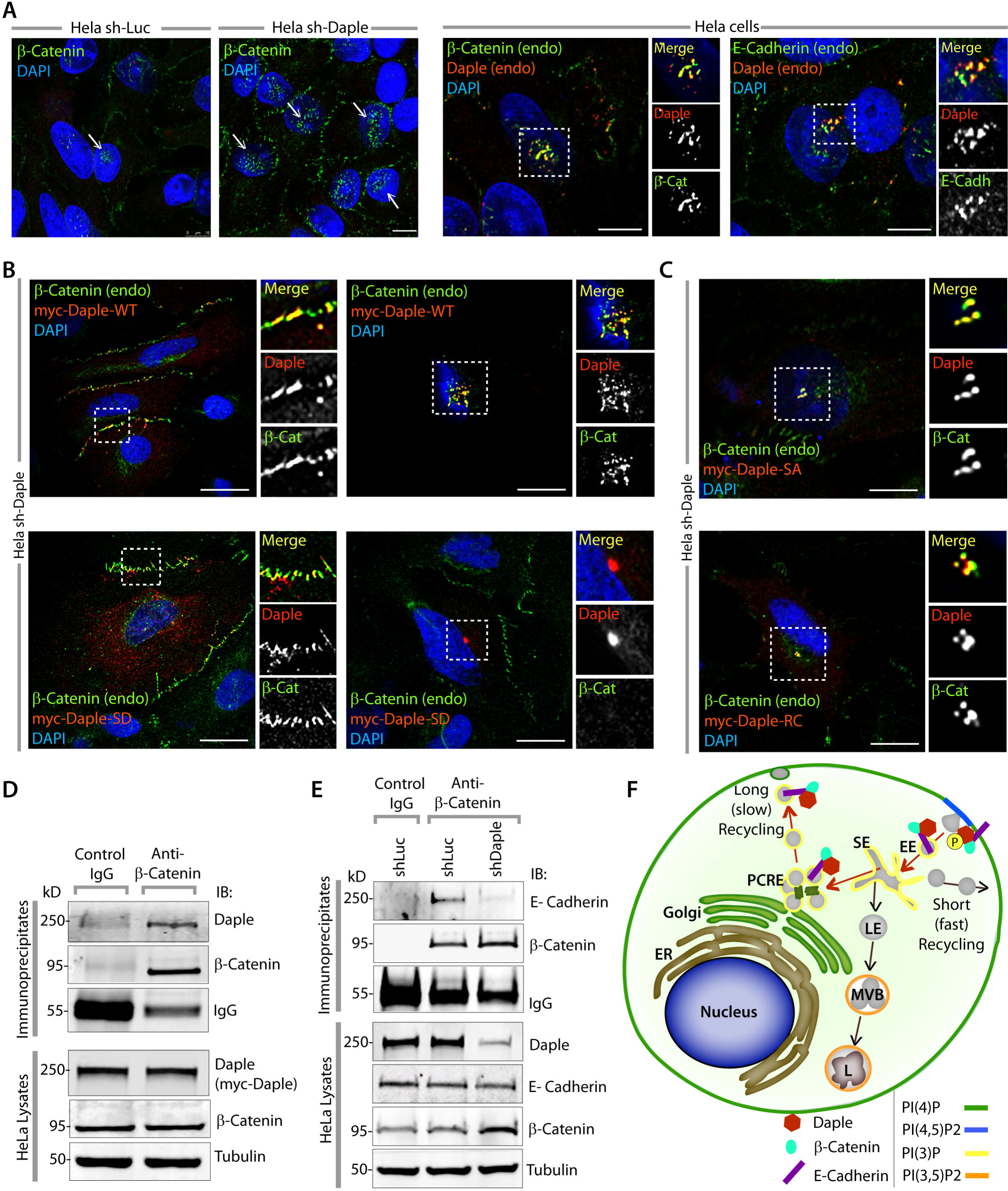
Daple binds β-catenin and regulates its distribution in cells. (**A**) <u>*Left*</u>: Control (shLuc) or Daple-depleted (shDaple) HeLa cells were analyzed for localization of β-catenin (Green) by immunofluorescence. <u>*Right*</u>: HeLa cells were analyzed for localization of endogenous Daple (red) and β-catenin or E-cadherin (Green) by immunofluorescence. Scale bar = 10 μm. (**B-C**) Phosphorylation of Daple on Ser1428 by Akt regulates the subcellular localization of β-catenin. Daple-depleted (shDaple) HeLa cells expressing myc-Daple-WT (B, top), myc-Daple SD (**B**, bottom), myc-Daple SA (**C**, top) and myc-Daple RC (**C**, bottom) were analyzed for localization of myc-Daple (red) and β-catenin (green) by confocal microscopy. Scale bar = 15 μm. (**D**) Daple interacts with β-catenin in cells. β-catenin was immunoprecipitated from equal aliquots of lysates (bottom) of HeLa cells expressing myc-Daple-WT using anti-β-catenin mAb or control IgG. Immune complexes (top) were analyzed for Daple, β-catenin and Tubulin by immunoblotting (IB). (**E**) β-catenin was immunoprecipitated from equal amounts of lysates of control (shLuc) and Daple-depleted (shDaple) HeLa cells as in **D** and immune complexes were analysed for E-cadherin by immunoblotting (IB). E-cadherin was detected in the β-catenin-bound complexes in the presence of Daple, whereas it was virtually undetectable in shDaple cells (right lane) and in control IgG (left lane). (**F**) Working Model: Schematic summarizing the trafficking itinerary (red arrows) of Daple-bound β-catenin-E-cadherin complexes from cell-cell contact sites at the PM and the pericentriolar recycling endosomes [PCREs]. The distribution of β-catenin is regulated by Daple’s ability to reversibly bind phosphoinositide PI3P on endosomes, which in turn is regulated by phosphorylation of S1428 on Daple by Akt. In the absence of phosphorylation (SA and RC mutants), Daple constitutively remains bound to PI3-P, and is found predominantly at the PCREs. When constitutively phosphorylated (SD), β-catenin and Daple preferentially colocalize at cell-cell contacts. Key: EE, early endosomes; SE, sorting endosomes; LE, late endosomes; PCRE, peri-centriolar recycling endosomes; ER, endoplasmic reticulum; MVB, multi-vesicular bodies; L, lysosomes.

First, we confirmed that Daple colocalizes with β-catenin and E-cadherin in the pericentriolar compartment (**Figure 4A; right**), as determined by confocal immunofluorescence analyses on endogenous proteins in HeLa cells. Next, we transfected Daple-WT or mutants in HeLa cells depleted of endogenous Daple, and looked for β-catenin distribution in cells. We found that two distinct patterns of localization were seen for Daple-WT at equal frequency, i.e., predominantly at the PM in ~ 50% cells (**Figure 4B**; top left) or predominantly within PCREs in ~ 50% cells (**Figure 4B**; top right); Daple colocalized with β-catenin at both locations. In the case of Daple-SD, this mutant also showed two patterns of localization, at the PM and cell-cell contact sites (~ 80%) and in PCREs (~ 20%). Unlike Daple-WT, Daple-SD colocalized with β-catenin exclusively at the PM (**Figure 4B**; bottom left) in 100% of the cells; no β-catenin was found in the PCREs (**Figure 4B**; bottom right). In the case of the non-phorphorylatable SA and RC mutants, Daple was found exclusively in PCREs (~ 100% of the cells), where it colocalized with β-catenin (**Figure 4C**); neither Daple, nor β-catenin was found at the PM or at cell-cell contact sites. These findings reveal a pattern—1) in cells expressing Daple-WT, β-catenin is distributed between 2 compartments at equal frequency, the PCREs and the cell-cell contact sites at the PM; 2) in cells expressing the phosphomimicking Daple-SD mutant, the distribution of β-catenin is skewed, i.e., shifted to the PM and cell-cell contacts, suggesting a failure to undergo endocytosis and reach the PCREs; 3) in cells expressing the non-phosphorylatable Daple-RC mutant, the distribution of β-catenin is skewed again, i.e., shifted to the PCREs, suggesting either a failure to be retained at the PM (i.e., rapidly endocytosed) or a failure to exit the PCREs and get exocytosed to the cell-cell contact sites at the PM. We also found that Daple and β-catenin interact in cells, as determined by coimmunoprecipitation of endogenous proteins from HeLa cells (**Figure 4D**), suggesting that these proteins likely interact within the macrocomplexes. We also confirmed that excess of β-catenin that accumulates in the expanded PCRE compartment of Daple-depleted HeLa cells (**Figure 4A**; left) is not complexed with E-cadherin, as determined by coimmunoprecipitation of endogenous Ecadherin/β-catenin complexes (**Figure 4E**). Because both phosphomimicking and nonphosphorylatable mutants distort the subcellular distribution of β-catenin, we conclude that reversible phosphoregulation of Daple’s PI3-P binding function by Akt is essential for normal co-distribution of Daple/ β-catenin/E-cadherin complexes in cells (see working model; **Figure 4F**). Phosphorylated Daple preferentially localizes at the PM and compartmentalizes β-catenin to cell-cell contact sites. Dephosphorylation is necessary to trigger endocytic trafficking of β-catenin from the PM to PCREs, likely because only dephosphorylated Daple can then engage with PI3-P on endosomes that enter the long-recycling pathway laden with these complexes. Daple that is constitutively dephosphorylated preferentially localizes to the PCREs and compartmentalizes β-catenin in that location.

### Daple binds and colocalizes with dynein at PCREs; phosphorylation of Daple by Akt alters dynein localization

The co-distribution of Daple/ β-catenin/E-cadherin complexes at the PM and on PCREs raises the possibility that directed endocytic traffic of these complexes over long-distances between these compartments must use motor proteins. In fact, prior studies have shown that during the endocytosis of β-catenin/E-cadherin complexes, clathrin/AP-2 is released from endocytic vesicles, allowing dynein recruitment for retrograde transport (Skanland et al, 2009); dynein directly binds β-catenin, captures and tethers microtubules at cell-cell contact sites to initiate long-distance transport (Ligon & Holzbaur, 2007; Ligon et al, 2001). We found that both endogenous and exogenously expressed Daple colocalizes with dynein on PCREs (**Figure 5A; S4A**); such colocalization is preserved in the case of Daple-FA and Daple-ΔPBM (**Figure 5A**), indicating that Daple colocalizes with dynein regardless of its ability to bind or activate G proteins or bind Dvl. Coimmunoprecipitation studies confirmed that endogenous Daple and dynein interact (**Figure 5B**). Exogenous expression of Daple in either control (shLuc) or Daple-depleted HeLa cells resulted in a robust enrichment of endogenous dynein at PCREs (**Figure S4B**), suggesting that the interaction between Daple and dynein may enhance the activity of dynein. In wake of these findings, we hypothesized, that phosphoregulation of Daple by Akt may impact the localization of dynein/dynactin complex, and could account for the failure of endocytic trafficking of β-catenin from the PM to the PCREs in cells expressing Daple-SD. We found that is indeed the case, because while Daple-WT (**Figure 5C**), and both the non-phosphorylatable SA (**Figure 5D**) and RC (**Figure 5E**) mutants displayed robust enrichment of endogenous dynein in the PCREs, but Daple-SD did not (**Figure 5F-G**). Cells expressing Daple-SD formed cell-cell contacts frequently, where Daple-SD and dynein colocalized (**Figure 5F**). In cells that did not make contacts, Daple-SD was localized in vesicular structures in the cell periphery and in a pericentriolar compartment (likely PCREs), but dynein did not colocalize with Daple in these vesicles (**Figure 5G**). What may be the significance of Daple’s ability to bind and regulate the distributions of β-catenin and dynein in PCREs? We have confirmed that the PI3-P binding domain we identify here is not required for binding to dynein (unpublished data). Because Hook-domain containing proteins, Hook1 and 3 serve as adaptors that enhance the assembly and activation of dynein motor complex (Schroeder & Vale, 2016) to support unidirectional, long-range processive motility at high velocity (Olenick et al, 2016), it is possible that Daple, which has a Hook-domain at its N-terminus, also does the same. In fact, HkRP3, a paralogue of Daple has already been shown to bind dynein/dynactin complex and trigger the transport of lytic granules in cytotoxic natural killer (NK)-cells (Ham et al, 2015); whether Daple serves as a specific adaptor for dynein motor proteins for shuttling of β-catenin/E-cadherin cargo proteins remains to be seen.

**Figure 5:**
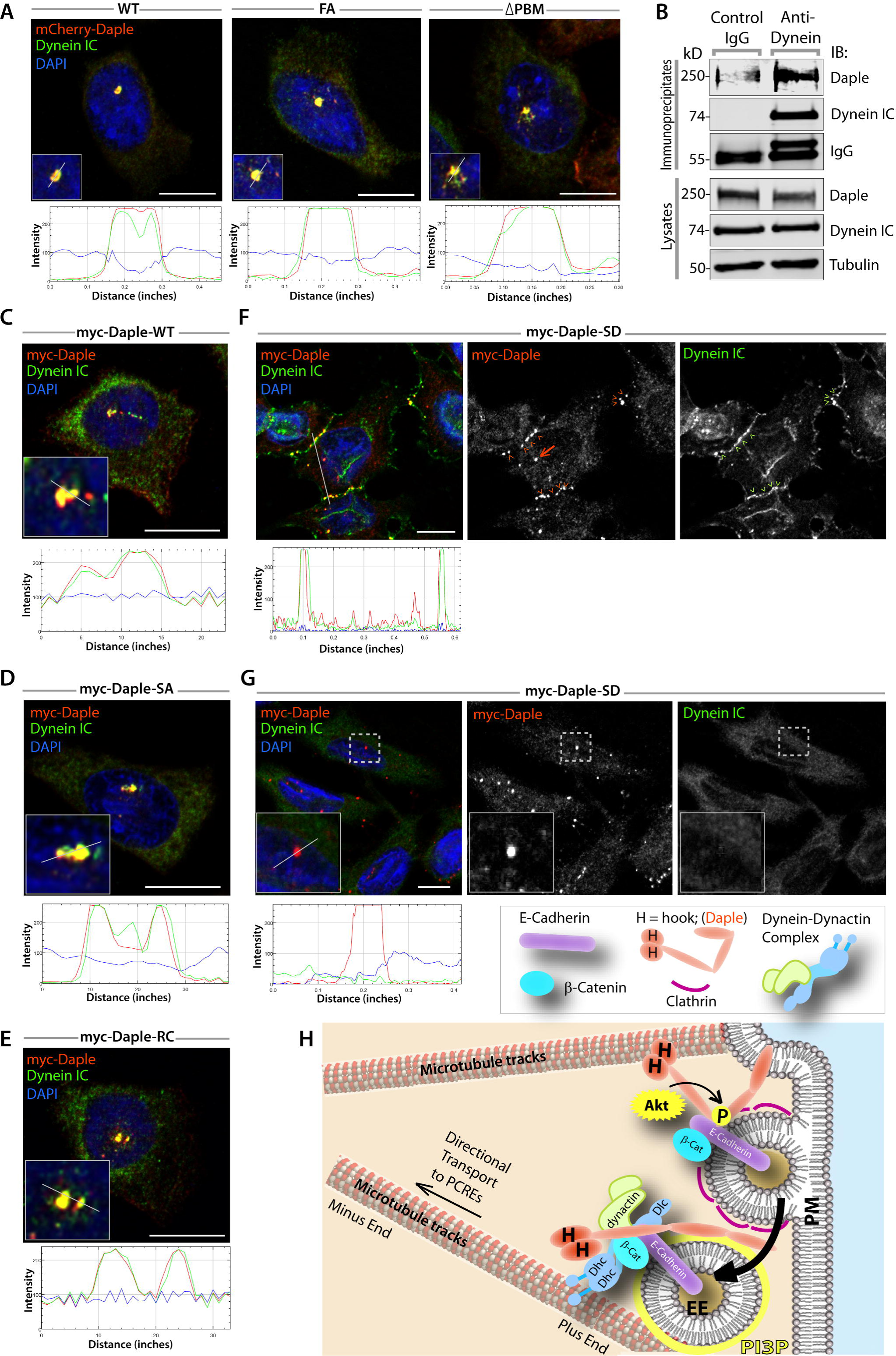
Daple binds and localizes with dynein in PCREs; phosphorylation at S1428 by Akt abolishes such colocalization. (**A**) HeLa cells expressing mCherry-Daple-WT (left), FA (middle; defective in binding G proteins), or ΔPBM (right; defective in binding Dvl) mutants (red) were analysed for endogenous dynein (intermediate chain, IC; green) and DAPI (nucleus) by confocal immunofluorescence. RGB plot, generated using ImageJ displayed underneath each image shows the degree of colocalization between red (Daple) and green (dynein) along the white line drawn in inset. Scale bar = 10 μm. (**B**) Dynein was immunoprecipitated from lysates of HeLa cells using control or anti-Dynein IC antibodies. Immunoprocipitates were analyzed for Daple and dynein IC by immunoblotting (IB). (**C-E**) Daple-depleted HeLa cells expressing myc-Daple-WT (**C**), SA (**D**), and RC (**E**) mutants were fixed, stained and analyzed for colocalization between myc-Daple (red) and endogenous dynein (green) by confocal immunofluorescence. RGB plots, generated using ImageJ displayed underneath each image shows the degree of colocalization between red (Daple) and green (dynein) along the white line drawn in inset. (**F-G**) Daple-depleted HeLa cells expressing myc-Daple-SD were fixed, stained and analyzed for the distribution of Daple (red) and dynein (green) by confocal imaging. Two patterns were noted: among cells with cell-cell contact (**F**), Daple and dynein colocalized primarily at the contact sites (arrowhead); a small pool of Daple was also seen at PCREs (arrow). Among cells that were not in contact, Daple-SD localized to PCREs and on peripheral vesicles, but dynein was not enriched at these sites. Scale bar = 10 μm. (**H**) Proposed working model for the role of the newly identified Akt→ phospho-Daple pathway in the distribution of β-catenin/Ecadherin complexes between the PM and the PCREs.

Based on our findings and the existing literature, we propose the following working model (**Figure 5H**): During the endocytosis of β-catenin/E-cadherin complexes, dynein/dynactin complexes are recruited to the β-catenin/E-cadherin complexes via a direct interaction with β-catenin (Ligon et al, 2001). Daple binds β-catenin and dynein, and colocalizes with β-catenin/E-cadherin complexes at cell-cell contact sites. PI3-K/Akt signaling that is initiated either by growth factors (Aznar et al., 2017; accompanying manuscript), or by Wnt5A/FZD7 (Aznar et al, 2015), or by E-cadherin within junctional complexes (Pece et al, 1999) phosphorylates Daple at S1428; phosphorylation helps Daple maintain β-catenin/E-cadherin complexes at cell-cell contacts, inhibit endocytosis of complexes, and thereby, facilitates adhesion. Dephosphorylation is required for Daple to bind PI3-P and facilitate endocytic trafficking of Daple-bound macrocomplexes [dynein/ β-catenin/E-cadherin] via the long-recycling pathway through Rab11-positive PCREs. Dephosphorylation also allows Daple to trigger the activation of dynein motor complex (Olenick et al, 2016) and movement of vesicles to minus ends of microtubules via mechanisms that currently remain poorly understood.

### Phosphorylation of Daple by Akt suppresses the canonical Wnt/ β-catenin pathway and growth in colonies, but enhances Rac1 activity, triggers migration and EMT

We previously demonstrated that Daple suppresses proliferation but enhances cell migration via activation of the non-canonical Wnt5A/FZD7 pathway (Aznar et al, 2015); it also does the same downstream of growth factor RTKs such as EGFR (Aznar et al., 2017; accompanying manuscript). To study the impact of phosphorylation of Daple at S1428 on those phenotypes, we generated HeLa cell lines stably depleted of endogenous Daple and rescued by expressing shRNA-resistant Daple-WT or various mutants at close to endogenous levels (**Figure 6A**). Rac1 activity, as determined in pulldown assays using the p21 binding domain (PBD) of PAK1, was enhanced in cells expressing the phorphomimicking SD mutant, and suppressed in those expressing the non-phosphorylatable SA and RC mutants (**Figure 6B-C**). These changes in Rac1 activity was accompanied also by changes in chemotactic migration of cells, as determined by transwell chemotaxis assays along a 0- 10% serum gradient (**Figure 6D-E**); compared to cells expressing Daple-WT, cells expressing the phosphomimicking Daple-SD migrated more, whereas those expressing the non-phosphorylatable SA or RC mutants migrated less. We conclude that phosphorylation of Daple at S1428 by Akt is required for enhancing Rac1 activity and chemotactic cell migration. The effect of this phosphorylation was reversed when it came to either anchorage-independent (**Figure 6F, G; S4B**) or anchorage-dependent (**Figure 6J, K, S4C**) colony formation; Daple-SA or RC mutants enhanced colony growth, whereas the Daple-SD mutant suppressed anchorage-dependent and virtually abolished anchorage independent colony growth.

**Figure 6:**
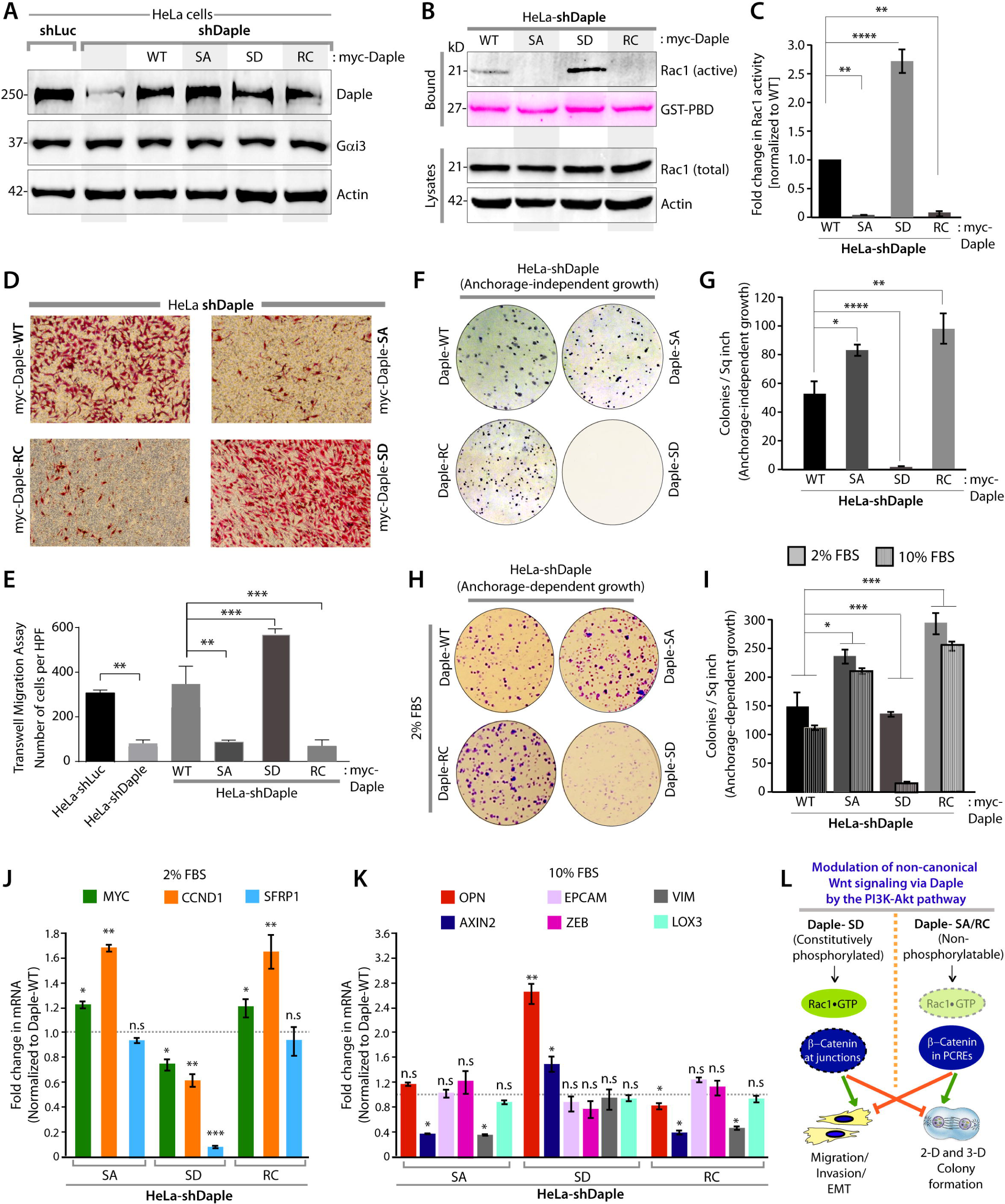
Constitutive phosphoactivation of Daple [mimicked by S1428D mutation] suppresses both anchorage-dependent and independent growth of tumor cell colonies, but enhances Rac1 activity and cell migration. (**A**) HeLa cell lines stably expressing various Daple constructs (see also *Methods)* were analysed for Daple and Gαi3 expression by immunoblotting. (**B**) Equal aliquots of lysates of HeLa cell lines were analyzed for active Rac1 using GST-PBD in pulldown assays, followed by immunoblotting. Compared to cells expressing Daple-WT, activation of Rac1 is impaired in cells expressing the non-phosphorylatable Daple SA and RC mutants, but enhanced in cells expressing the constitutively phosphomimicking SD mutant. (**C**) Bar graphs display the fold change in Rac1 activity. Error bars representing mean ± S.D of 3 independent experiments. (**D-E**) HeLa cell lines expressing various Daple constructs were analyzed for their ability to migrate in transwell assays toward a serum gradient (0.2% to 10% FBS). Images in **D** show representative fields of the transwell membrane, photographed at 60X mag. Compared to cells expressing Daple-WT, chemotactic migration is impaired in cells expressing the non-phosphorylatable Daple SA and RC mutants, but enhanced in cells expressing the constitutively phosphomimicking SD mutant. Graphs in **E** present the quantification of the number of migrating cells in **D**, averaged from 20 field-of view images per experiment (see also **Figure S4A**). Data are presented as mean ± SEM; n = 3. HPF = high-power field. (**F-I**) HeLa cell lines were analyzed for their ability to form colonies either in soft agar (F) or on plastic plates (**H**, 2% FBS; **S4C**, 10% FBS) for 2–3 weeks prior to fixing, staining, photography, and colony counting using an ImageJ Colony counter application (see also **Figure S4B-C** and *Methods*). Panel F shows representative photographs of the MTT-stained 6-well plates, whereas panel **J** shows representative photographs of colonies stained on plastic with crystal violet. Compared to cells expressing Daple WT, anchorage-independent (**F, G**) and anchorage-dependent (**H, I**) growth is impaired in cells expressing the constitutively phosphomimicking SD mutant, but enhanced in the cells expressing the non-phosphorylatable SA and RC mutants. Bar graphs in **G** and **I** display the number of colonies (y axis) seen in each cell line in **F, S4B, S4C** and **H**, respectively. \**p* < 0.05; \*\**p* < 0.01; \*\*\**p* < 0.001; \*\*\*\**p* < 0.0001. (**J-K**) HeLa cell lines in **A** were analyzed by qPCR for the levels of mRNA for the indicated canonical β-catenin/TCF/LEF target genes (**J**) or inducers of EMT (**K**). Results were normalized internally to mRNA levels of the housekeeping gene, GAPDH. Bar graphs display the fold change in each RNA (y axis) normalized to the expression in cells expressing Daple-WT (indicated with an interrupted grey line at 1.0). Error bars represent mean ± S.D of three independent experiments. \**p* < 0.05; \*\**p* < 0.01; \*\*\**p* < 0.001. (**L**) Schematic summarizing the impact of phosphorylation of Daple at S1428 on non-canonical Wnt signaling pathway and cellular phenotypes.

Prior studies by us (Aznar et al, 2015); Aznar et al., 2017; accompanying manuscript) and others (Ara et al, 2016; Ishida-Takagishi et al, 2012) have shown that non-canonical Wnt signaling that is enhanced by Daple displays a bifaceted transcriptional response: 1) it antagonistically suppresses canonical Wnt signaling and reduces the expression of β-catenin/TCF/LEF target genes; and 2) it enhances the expression of genes that trigger EMT. We found that constitutive phosphoactivation of Daple by Akt [mimicked by SD mutation] supresses several key canonical Wnt responsive β-catenin/TCF/LEF target genes (MYC, CCND1 and SFRP1; **Figure 6J**), consistent with its observed growth suppressive properties. Surprisingly, two other targets of β-catenin, osteopontin [OPN; which is known as a master regulator of EMT (Kothari et al, 2016) via its ability to stabilize vimentin (Dong et al, 2016)] and Axin2 [AXIN2, which enhances EMT via induction of Snail (Yook et al, 2006)] were not suppressed, and instead were increased in Daple-SD cells (**Figure 6K**), also consistent with its promigratory phenotype. Constitutive inactivation of the Akt→pS1428-Daple axis [as mimicked by SA and RC mutants] was associated with enhanced transcription of β-catenin/TCF/LEF target genes (MYC and CCND1; **Figure 6J**) and suppression of key inducers of EMT (AXIN2 and VIM; **Figure 6K**); all consistent with the pro-growth and poorly motile phenotype of these cells. It is noteworthy that all 3 cell lines expressing Daple mutants [SD, SA and RC] had elevated levels of β-catenin compared to cells expressing Daple-WT (**Figure S4D**), suggesting that the differential effects of phosphomimicking and non-phosphorylatable Daple mutants on the β-catenin/TCF/LEF axis is largely due to the differential localization of β-catenin. Selective enhancement of some EMT-associated β-catenin/TCF/LEF–dependent target genes (OPN and AXIN2) and suppression of other proliferation-associated transcriptional targets (MYC, CCND1 and SFRP1) in Daple-SD cells (**Figure 6J, K**), despite the relatively lower levels of β-catenin in these cells compared to SA/RC cells (**Figure S4D**) was an unexpected observation. Such differential response of β-catenin/TCF/LEF target genes in Daple-SD cells could be due to crosstalk with alternative pathways that can affect OPN and AXIN2, such as the Hippo signaling pathway (Lu et al, 2010; Park et al, 2015) or Rac1; the latter has been implicated in transcriptional upregulation of genes involved in EMT and cell invasion (Iwai et al, 2010; Jamieson et al, 2015; Wu et al, 2008).

Regardless of the mechanisms involved, our findings demonstrate that phosphorylation of Daple on residue Ser1428 is a key determinant of cellular phenotype (**Figure 6L**); a constitutively phosphorylated state in which β-catenin localizes to cell-cell contact sites, suppresses the β-catenin/TCF/LEF pathway, suppresses colony formation, but confers migratory advantage and triggers EMT. By contrast, a constitutively dephosphorylated state in which β-catenin predominantly localizes to PCREs, enhances key β-catenin/TCF/LEF target genes (MYC and CCND1), suppresses EMT, and thereby, confers a proliferative advantage over migration. We conclude that the skewed redistribution of β-catenin we see in phosphomimic (SD) or non-phosphorylatable (SA/RC) mutants may in part be responsible for the observed cellular phenotypes.

### Daple and Akt enhance non-canonical Wnt signaling within a feed-forward loop; implications in normal *vs*. transformed cells

Non-canonical Wnt signaling via Daple has been shown to serve as a double-edged sword; it serves as a tumor suppressive pathway in normal cells and early during tumorigenesis, but enhances EMT and metastatic progression later during cancer progression (Aznar et al, 2015). Based on our current findings, we propose the following model (**Figure 7A**): When phosphorylated by Akt, Daple’s ability to maintain large pools of β-catenin within Ecadherin/ β-catenin complexes at the cell-cell contact sites and enhance non-canonical Wnt signaling may primarily serve as a check and balance for the canonical Wnt pathway and to suppress proliferation. We previously showed that activation of Gαi proteins by Daple downstream of Wnt5A/FZD7 receptor enhances the PI3-K/Akt signaling pathway (Aznar et al, 2015). We hypothesized that the Akt→Daple axis we defined here may serve as a forward-feedback loop for the previously defined Daple→Akt axis. To test that, we analyzed Akt-mediated phosphorylation of Daple-WT or Daple-FA; the latter cannot activate G proteins and is defective in enhancing Akt signaling. We found that Akt phosphorylated Daple-WT ~2.5 fold more efficiently than it did Daple-FA (**Figure 7B**), indicating that the Akt↔Daple crosstalk is indeed bidirectional and capable of serving as a feedforward loop in cells.

**Figure 7:**
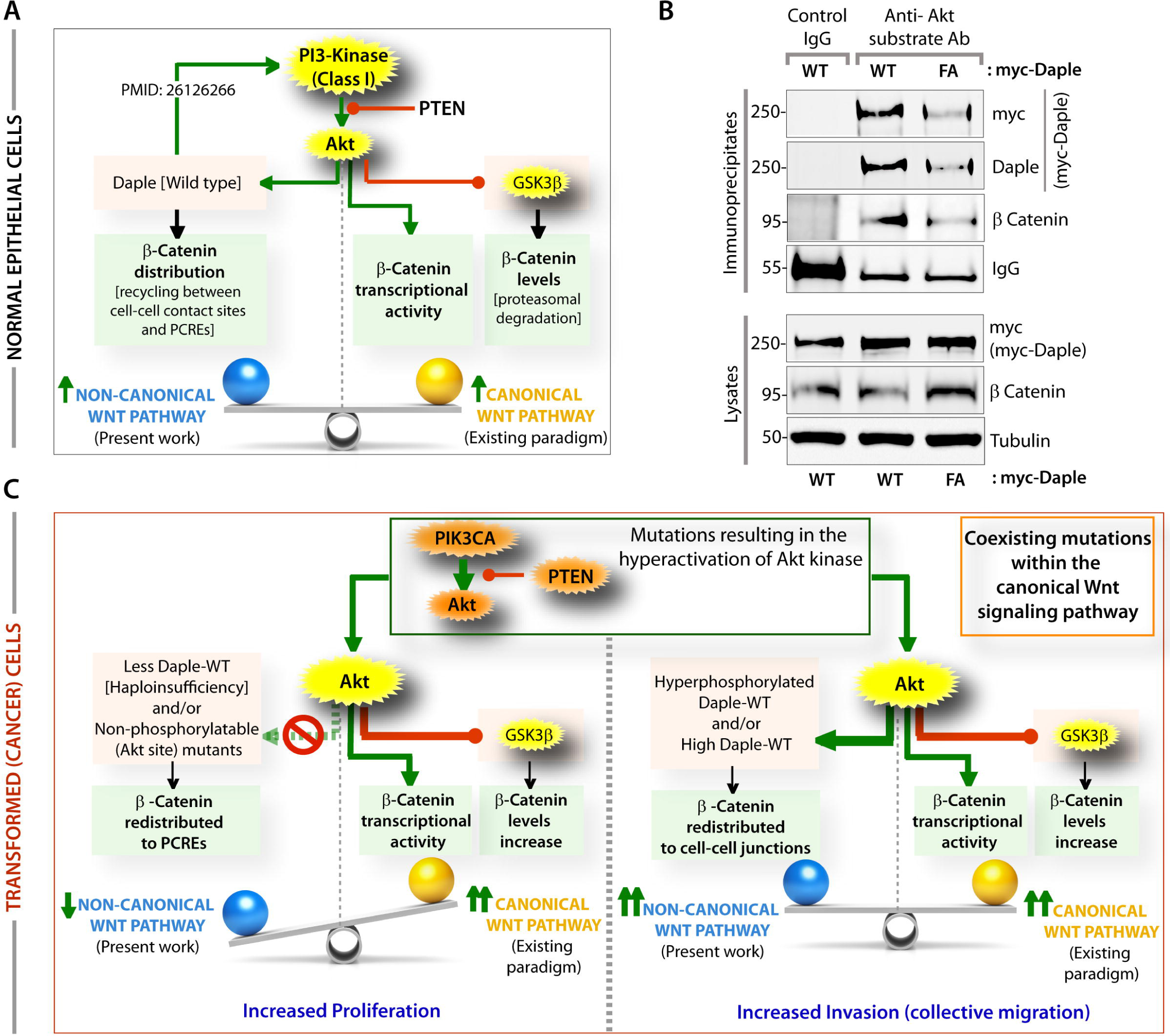
Phosphoregulation of Daple by Akt serves as a forward feedback loop for Daple’s GEF activity-dependent Akt signaling; this feed-forward loop regulates β-catenin distribution in cells. (**A**) Working model for Akt-Daple crosstalk in normal cells: Activation of the PI3K-Akt axis is known to enhance canonical Wnt signaling (right) via the upregulation of both transcriptional activity (Fang et al, 2007) and stability [inhibits proteasomal degradation] of β-catenin. Akt also phosphorylates Daple and enhances non-canonical Wnt signaling, regulates the intracellular distribution of β-catenin between the PCRE and the cell-cell junctions. Daple has previously been shown to enhance Akt phosphorylation (Aznar et al, 2015). (**B**) Daple was immunoprecipitated from equal aliquots of lysates of HeLa cells expressing myc-Daple [WT and FA] using anti-Akt-substrate antibody or control IgG. Immune complexes (top) were analyzed by immunoblotting (IB) for Daple using anti-Daple and anti-Myc. Compared to Daple WT, Akt-mediated phosphorylation of Daple was reduced in Daple FA. (**C**) Working model for Akt-Daple crosstalk in transformed cells: Mutations commonly affect the PI3K-Akt pathway in tumor cells, causing an overall increase in Akt signaling. Under these circumstances, canonical Wnt signaling is enhanced via mechanisms mentioned in **A**. As for non-canonical Wnt pathway, the effect of aberrant Akt signaling varies depending on the levels of expression or mutations in Daple. <u>*Left*</u>: If Daple is low or is expressed as a non-phosphorylatable mutant, non-canonical Wnt signals are suppressed. <u>*Right*</u>: If Daple is high and expressed as a WT protein, it can get hyperphosphorylated by Akt, in which case, non-canonical Wnt signaling is enhanced.

As for transformed cancer cells, which accumulate multiple genetic anomalies, frequently involving constitutive hyperactivation of the PI3-K/Akt cascade (e.g., activating mutations in PIK3CA or Akt and inactivating mutations in PTEN) and within the canonical Wnt/ β-catenin signaling axis (e.g., APC, Axin, β-catenin, or GSK3β), the impact of the Akt↔Daple crosstalk may fuel cancer initiation and/or progression. When Daple is low or the Akt→Daple axis is disrupted, as in tumors harboring Daple-R1425C or S1428F mutations (**Figure 2B**), β-catenin cannot be retained at cell-cell contact sites within E-cadherin/ β-catenin complexes, and instead reaches the PCREs. Consequently, non-canonical Wnt signals cannot be enhanced and canonical β-catenin-dependent Wnt signaling goes unchecked. When Daple levels are high and it is hyperphosphorylated by Akt, a large pool of β-catenin is maintained within E-cadherin/ β-catenin complexes at the cell-cell contact sites, non-canonical Wnt pathways (Rac1 and EMT) are triggered, while maintaining cell adhesion, perhaps enabling collective tumor cell migration during cancer invasion. This form of migration, in which groups of cells invade the peritumoral stroma while maintaining cell–cell contacts, is the predominant form of migration displayed in invasive solid tumours (Clark & Vignjevic, 2015; Friedl et al, 2012) and involves activation of Rac1 and PI3-K/Akt signals (Yamaguchi et al, 2015); two pathways that are hallmark of Daple-dependent signaling (Aznar et al, 2015).

## Conclusions

The major finding in this work is the discovery of a novel crosstalk between the PI3-K/Akt and the non-canonical Wnt pathways via Daple which results in compartmentalization of β-catenin. Widely assumed as an enhancer of the canonical Wnt signaling pathway, Akt has been implicated in enhancing the TCF/LEF transcription machinery by either directly phosphorylating β-catenin (Fang et al, 2007), or by inhibiting GSK3β, a negative regulator of β-catenin stability (Aberle et al, 1997; Salic et al, 2000). Here we show how a single phosphoevent triggered by Akt targets Daple’s PI-binding domain and imparts spatial regulation of non-canonical signaling via Daple. In doing so, Akt directly impacts Daple’s ability to associate with PI3-P-enriched endosomes that engage dynein motor complex for long-distance travel to PCREs. These PCREs carry β-catenin/E-cadherin complexes as cargo proteins whose localization is impacted by spatially regulated signaling via Daple. Phosphorylation by Akt compartmentalizes Daple and β-catenin/E-cadherin at cell-cell contact sites, which suppresses the canonical β-catenin/TCF/LEF transcriptional machinery and thereby, suppresses growth. These findings are in agreement with prior work showing that the tumor suppressive effects of non-canonical Wnt signaling is due, in part, to its ability to sequester β-catenin at the PM (Bernard et al, 2008). It is also in keeping with the widely-accepted role of β-catenin/E-cadherin complexes at the cell junctions in the maintenance of cell polarity, which is a gatekeeper against cell proliferation and cancer initiation (Royer & Lu, 2011). We conclude that compartmentalization of β-catenin at cell-cell contact sites inhibits proliferation in part by prolonging non-canonical Wnt signals and tonic suppression of canonical Wnt signals. When the Akt→Daple axis is disrupted, both Daple and β-catenin cannot be maintained at cell-cell contact sites, and instead, they are endocytosed and trafficked to PCREs; consequently, the non-canonical Wnt signals are inhibited, canonical Wnt pathway goes unopposed, and cells proliferate. That Daple/β-catenin/E-cadherin/dynein complexes compartmentalize to PCREs is not surprising because prior work has documented the presence of multiple components of the Wnt signalosome on PCREs [e.g., β-catenin/Amotl2 complexes (Li et al, 2012), Dvl, (Smalley et al, 2005), FZD7 (Egea-Jimenez et al, 2016; Pradhan-Sundd & Verheyen, 2015)] and the role of endocytosis and recycling via Rab11 positive PCREs in the regulation of the Wnt signalosome (Feng & Gao, 2015), and specifically, in the termination of non-canonical Wnt signals (Gagliardi et al, 2008). Because canonical signal transduction (via β-catenin stabilization) requires endocytosis, and endosomes are believed to serve as critical platforms for spatially restricted canonical Wnt signaling (Disanza et al, 2009), we conclude that compartmentalization of β-catenin in PCREs triggers proliferation in part by prolonging unopposed canonical Wnt signaling.

It is noteworthy that all findings reported here were observed at steady-state without subjecting cells to acute stimulation with ligands, largely because Daple’s localization to the PCREs appeared to be constitutive and independent of its ability to bind G proteins or Dvl, two partners that have been shown to be critical for Daple’s role in Wnt5A/FZD7 (Aznar et al, 2015) and growth factor RTK (Aznar et al., accompanying manuscript, 2017) signaling. Which begs the question, that in the absence of acute ligand stimulation, or activating mutations, what other alternative pathways for Akt activation is responsible for the observed effects. Prior studies have shown that much like FZD7 and RTKs, E-cadherin/β-catenin complexes are capable of initiating outside-in PI3-K/Akt signaling that is critical for aggregation-dependent cell survival (Pece et al, 1999); E-cadherins at cell-cell contact sites trigger rapid PI3-K-dependent activation of Akt. It is possible that the Akt→Daple→β-catenin redistribution pathway we define here is initiated by E-cadherins within E-cadherin/ β-catenin complexes at cell-cell contact sites.

In conclusion, we provide the first direct evidence here that Akt can not only enhance the canonical Wnt pathway, but also enhance the non-canonical Wnt pathway by phosphorylating Daple. Insights gained also reveal how this crosstalk in the setting of aberrant activation of the PI3-K/Akt pathway impacts cancer initiation and/or progression by modulating both Wnt pathways.

## Materials and Methods

*Detailed Methods can be found in Supplementary Information*.

### Protein Expression and Purification

GST and His-tagged recombinant proteins were expressed in E. coli strain BL21 (DE3) (Invitrogen) and purified as described previously (Garcia-Marcos et al, 2011; Ghosh et al, 2010; Ghosh et al, 2008).

### Transfection; Generation of Stable Cell Lines and Cell Lysis

Transfection was carried out using either polyethylenimine (PEI) as previously described (Aznar et al, 2016) or Genejuice from Novagen for DNA plasmids following the manufacturers’ protocols. Selection or stable cells, triton extracts and whole cell lysis were carried out as mentioned before (Aznar et al, 2015).

### Quantitative Immunoblotting

Infrared imaging with two-color detection and band densitometry quantifications were performed using a Li-Cor Odyssey imaging system exactly as done previously (Aznar et al, 2015). All Odyssey images were processed using Image J software (NIH) and assembled into figure panels using Photoshop and Illustrator software (Adobe).

### In vitro GST pulldown and Immunoprecipitation Assays

These assays were carried out exactly as previously (Aznar et al, 2015) and are detailed in *Supplementary Information*.

### In vitro and in-cellulo Kinase Assays

In vitro kinase assays were performed using bacterially expressed His (6 x His, hexahistidine) tagged Daple (aa 1194-1491) proteins (~3-5 µg per reaction), and ~50 ng recombinant kinases which were obtained commercially (SignalChem, Canada). The reactions were started by addition of ATP (5 µCi/reaction vP32-ATP and 6◻µM cold ATP) and carried out at 30°C in 30◻µl of kinase buffer (20◻mM Tris·HCl, pH 7.5, 2◻mM EDTA, 10◻mM MgCl2 and 1◻mM DTT). Phosphorylated proteins were separated on 10% SDS-PAGE and detected by autoradiography.

For *in cellulo* phosphorylation assays anti-phosphoAkt substrate Ab was used to immunoprecipitated phosphoproteins from cell lysates [prepared using RIPA buffer], and analysing the precipitates for Daple by immunoblotting.

### Lipid blot assay

The lipid blots were performed as previously described (Dippold et al, 2009). PVDF membranes spotted with lipids were incubated with *in vitro* translated ^35^S-labeled Daple (aa 1194-1491).

### Rac1 activity, transwell migration, colony growth, and qPCR assays

These assays were carried out exactly as outlined before (Aznar et al, 2015).

### Statistical analysis

Each experiment presented in the figures is representative of at least three independent experiments. Displayed images and immunoblots are representative of the biological repeats. Statistical significance between the differences of means was calculated by an unpaired student’s t-test or one-way ANOVA [whenever more than two groups were compared]. A two-tailed *p* value of <0.05 at 95% confidence interval is considered statistically significant. All graphical data presented were prepared using GraphPad or Matlab.

## ACKNOWLEDGMENTS

We thank Gordon N. Gill and Marilyn G. Farquhar (UCSD) for their critical input during the preparation of the manuscript. This work was supported by NIH grants CA100768, CA160911 and DK099226 (to P.G). P.G. was also supported by the UC San Diego Moores Cancer Center. MDB is funded by the American Cancer Society by the Lee National Denim Day Postdoctoral Fellowship Award PF-13-367-01-CDD.

## AUTHOR CONTRIBUTIONS

N.A performed and analyzed most of the experiments in this work. N.S carried out the GST-FYVE binding assays. Y.D carried out the qPCR assays. Y.D and J.E cloned and generated all Daple mutants used in this work. M.B and S.J.F carried out the lipid blots. N.A and P.G conceived the project and wrote the manuscript. P.G supervised and funded the project.

